# Eastern equine encephalitis virus-vaccinated mice are protected against Madariaga virus despite negligible neutralizing antibody titers

**DOI:** 10.64898/2026.09.14.751423

**Authors:** Samuel A. Cerezo, Chi-Hsuan Sung, Wendy W. Tang, Gabriel L. Hamer, Raquel R. Rech, Tereza Magalhaes

**Affiliations:** Department of Entomology, Texas A&M University, College Station, TX, USA; Department of Veterinary Pathobiology, Texas A&M University, College Station, TX, USA

**Keywords:** Alphavirus, *Alphavirus madariaga*, cross-protection, arbovirus, equine encephalitides, inactivated vaccines, heterologous immunity

## Abstract

Madariaga virus (MADV) is an understudied member of the eastern equine encephalitis virus (EEEV) complex that circulates widely in Latin America and may be geographically expanding. Spillover infections can cause severe disease in humans and equids. Despite these public and animal health concerns, no MADV vaccines or treatments are available. In this context, commercially available equine vaccines against the closely related North American EEEV (NA EEEV) could potentially provide heterologous protection against MADV. However, previous studies have shown weak or undetectable MADV-neutralizing antibody responses in equids and humans immunized with inactivated NA EEEV vaccines, and whether protection can occur despite these limited neutralizing antibody responses remains unknown. We evaluated a commercial equine trivalent inactivated NA EEEV vaccine against MADV challenge in NIH Swiss mice. Animals received two vaccine doses 14 days apart and were challenged 14 days later. Vaccination provided protection against lethal disease, whereas 36% mortality occurred in the unvaccinated group. Overall, vaccinated animals developed fewer and less severe clinical signs. MADV RNAemia was not detected following vaccination, and tissue dissemination was significantly reduced compared with the unvaccinated group. Central nervous system lesions were observed only in the unvaccinated group. Despite protection, most vaccinated mice had no detectable MADV-neutralizing antibody titers, and those that seroconverted developed only low titers, whereas most unvaccinated mice developed intermediate to high titers. These findings demonstrate that an inactivated NA EEEV-containing vaccine can protect against severe MADV disease despite limited MADV-neutralizing antibody responses.

**IMPORTANCE:** Madariaga virus (MADV) is an emerging mosquito-borne alphavirus that can cause severe disease in humans and equids and may be expanding its geographic range in the Americas. Yet no vaccines or treatments targeting MADV are currently available. We show that an existing commercial equine vaccine against the related North American eastern equine encephalitis virus protected mice against severe disease and death following MADV exposure. Importantly, protection occurred even though most vaccinated animals had no detectable MADV-neutralizing antibodies, while a few had low titers. These findings raise the possibility that existing equine vaccines could provide protection during MADV outbreaks and highlight the need to determine whether similar protection occurs in equids. They also indicate that immune responses other than neutralizing antibodies may contribute to protection and should be considered when evaluating immunity elicited by inactivated vaccines.

## INTRODUCTION

Madariaga virus (MADV), which comprises Lineages II–IV of the eastern equine encephalitis virus (EEEV) complex, is a mosquito-borne alphavirus that circulates widely in Latin America (1). North American EEEV (NA EEEV) constitutes Lineage I of the complex (1, 2). Although its mosquito vectors have not been definitively identified, field and laboratory evidence implicates *Culex* (*Melanoconion*) spp. as potential enzootic vectors and *Aedes taeniorhynchus* as a putative epizootic vector (1, 3–6). The transmission cycle of MADV appears more similar to that of Venezuelan equine encephalitis virus (VEEV) than that of NA EEEV, with small ground-dwelling mammals, such as rodents, suggested as vertebrate reservoirs (2).

MADV is as pathogenic to equids as NA EEEV, with case fatality rates ranging from 40 to >90% during equine epizootics in Latin America (1). In humans, MADV has historically been considered less virulent than NA EEEV because fatal cases have rarely been reported (1, 7, 8). However, an outbreak of human encephalitis in Panama in 2010 challenged this paradigm and demonstrated the virus’ potential to cause outbreaks involving severe neurological disease (9). In addition, MADV has been detected, in some instances incidentally, in individuals presenting with acute febrile illness in Haiti and other Latin American countries (10–12). Limited recognition of MADV as a cause of human disease, together with clinical manifestations that may overlap with those of other arboviral infections, could contribute to underrecognition and underreporting of human cases, particularly in areas where arboviruses such as dengue and chikungunya viruses are endemic. The detection of MADV in Haiti, where the virus had not previously been reported, may also indicate ongoing geographic expansion (12).

MADV and NA EEEV are genetically related but antigenically distinct, sharing approximately 23-24% nucleotide divergence and 9-11% amino acid divergence within the structural polyprotein (2). Historically classified as the South and North American subtypes within the EEEV complex, respectively (13), they are now recognized as separate species, with MADV classified as *Alphavirus madariaga* (14).

There are no approved vaccines against MADV for humans or equids. For NA EEEV, an inactivated human vaccine (PE-6) was previously available under an Investigational New Drug (IND) permit for military and laboratory personnel at risk (15–17), while several multivalent inactivated vaccines are licensed for use in equids (18). Immunity elicited by the equine vaccines is generally considered short-lived, and annual booster vaccinations are therefore recommended (19).

In the absence of an approved MADV vaccine, existing NA EEEV vaccines could provide an alternative strategy for the prevention and control of MADV outbreaks. Evidence of cross-protection would indicate an added benefit of NA EEEV vaccination in regions where MADV already circulates and could support the deployment of these vaccines should MADV emerge in new regions. However, the limited available evidence indicates weak heterologous immunity between NA EEEV and MADV based on neutralizing antibody titers. For instance, during a MADV equine epizootic in Panama in 1973, surviving animals with high neutralizing antibody titers against MADV had low titers against NA EEEV, suggesting limited heterologous neutralizing antibody responses following natural infection (20). Similarly, only 33% of horses vaccinated with an inactivated bivalent EEEV/western equine encephalitis virus (WEEV) vaccine developed detectable neutralizing antibodies against MADV, compared with 75% against NA EEEV (20). Limited heterologous neutralizing antibody responses have also been observed in humans, with volunteers vaccinated with the inactivated PE-6 NA EEEV vaccine developing undetectable or low titers against MADV, leading the authors to conclude that vaccination against NA EEEV was unlikely to confer protection against MADV (16).

Whether the low heterologous neutralizing antibody responses observed following NA EEEV vaccination translate into limited protection against MADV remains unknown. We therefore evaluated the protective efficacy of a trivalent inactivated equine NA EEEV vaccine against MADV challenge in a mouse model. In addition, this study contributes to our understanding of the immune response elicited by the inactivated NA EEEV vaccine in the context of heterologous challenge.

## MATERIALS AND METHODS

### Virus and mice

The virus used in the experiments was MADV Brazil 165 (MADV-BR; GenBank: MZ389692.1), isolated by our group from a horse in northeastern Brazil that presented with clinical signs and died of neurological disease (21). For the challenge, a previously prepared frozen stock of passage 11 MADV-BR grown in Vero cells (ATCC CCL-81) with a titer of 1.2×10^7^ PFU/mL was used.

NIH Swiss mice were selected based on this strain’s susceptibility to MADV (22).

### Vaccine and vaccination strategy

Mice were vaccinated with Prestige 3® (Merck Animal Health, Intervet Inc., Omaha, NE, USA), a commercially available trivalent equine vaccine prepared with inactivated EEEV, inactivated WEEV, and tetanus toxoid. The vaccine also contains Havlogen® as an adjuvant. Prestige 3® comes in liquid form and the preconized vaccine dose for horses is 1 mL.

Fifty-four, 5-week-old NIH Swiss mice were acquired from Charles River Laboratory and acclimated for three days at the Texas A&M University (TAMU) Laboratory Animal Research and Resources facility. Mice were split into the following experimental groups: 1) Prestige 3® vaccination and mock (Dulbecco’s Modified Eagle Medium [DMEM]) challenge (Group 1); 2) Mock (Phosphate Buffered Saline [PBS]) vaccination and MADV challenge (Group 2); and 3) Prestige 3® vaccination and MADV challenge (Group 3). Group 1 initially consisted of 5 males and 5 females, but one female died before challenge following an unrelated handling event. Groups 2 and 3 consisted of 11 females and 11 males each. The smaller sample size of Group 1 was by design, as this group was originally included as a control for pathology analysis only; however, Group 1 animals were subsequently included in the clinical data analysis. All mice had access to food pellets and water ad libitum.

Mice received two doses of vaccine fourteen days apart. Doses consisted of 200 µL of either vaccine or PBS. Fourteen days after the second dose (boost), mice were challenged with MADV-BR or mock challenged with DMEM. The challenge dose was 10,000 plaque forming units (PFUs) of MADV-BR diluted in DMEM (100 µL total volume) or 100 µL of DMEM only.

The experimental design is summarized in Fig. 1.

**Fig 1.**
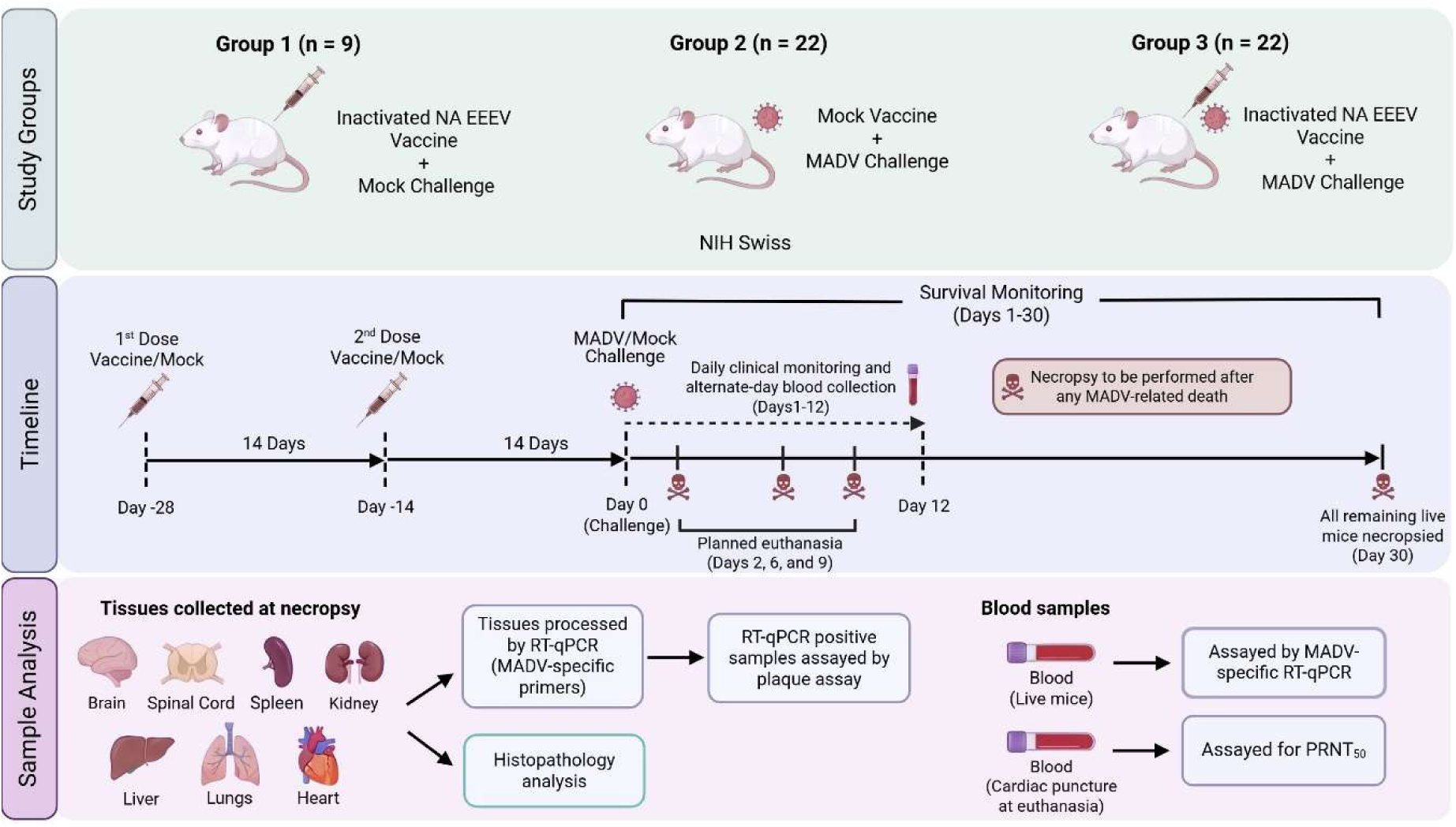
Experimental design and sample analysis workflow. Overview of the experimental groups, vaccination and challenge schedule, clinical monitoring and blood sampling timeline, planned euthanasia time points, and laboratory analyses. See Methods for detailed experimental procedures. Figure created in BioRender (Magalhaes, T. (2026); https://BioRender.com/rli40qw).

### Sample collection

Blood was collected one day before the first dose of vaccination, one day before the second dose, one day before challenge, and on alternating days throughout a 12-day period after challenge. Blood samples (collected through cheek or tail bleeds) were centrifuged (2,000xg, 10 min, 15°C) for serum separation, which was then transferred to clean microcentrifuge tubes. Sera were stored at −80°C until further use.

Mice were euthanized at predetermined time points for histopathologic analysis (2, 6, and 9 days post-inoculation [dpi]), upon reaching humane endpoints due to disease, or at the end of the study (30 dpi). Euthanasia was performed by CO_2_ inhalation. Following cessation of breathing and observation of ocular pallor, exsanguination via cardiac puncture was performed as a secondary physical method to ensure irreversibility. Necropsy was performed immediately following euthanasia, and samples of the brain, spinal cord, spleen, kidney, liver, lungs, and heart were collected. For histopathology, tissue samples were fixed in 10% neutral buffered formalin. For RNA extraction and plaque assays, fresh samples of all above mentioned tissues, except the spinal cord, were collected into microcentrifuge tubes containing 500 µL of tissue diluent (PBS supplemented with 20% fetal bovine serum [FBS] and 1x Gibco Antibiotic-Antimycotic [Thermo Fisher Scientific, Waltham, MA, USA]), and 3-4 borosilicate beads. Fresh tissue samples were stored at −80°C until further use.

A final blood sample was collected by cardiac puncture at euthanasia and processed as described above.

### Survival and clinical signs monitoring

Data collection instruments were designed using the Research Electronic Data Capture (REDCap) system (23, 24), hosted at TAMU. Clinical sign selection was based on mouse clinical signs associated with distress, pain, and neurological disease (25), as well as consultation with a laboratory animal TAMU veterinarian. Initially, the monitored clinical signs included ruffled fur, lethargy, convulsions, limb paralysis, and coma. During the study, daily observations of experimental mice led to the addition of hunched back and orbital tightening as additional indicators of pain or distress. Clinical signs were categorized as absent, minor presence, or significant presence, and monitored for 12 days after challenge. Body weight was measured at baseline and daily for 12 dpi. Body temperature was initially measured using a digital thermometer, but because the measurements were found to be unreliable, temperature monitoring was discontinued.

Survival was monitored for 30 days to detect potential delayed disease progression. Mice that reached the humane endpoints due to diseases defined in the approved animal protocol were euthanized.

### Viral RNA and infectious virus load in tissues

To assess the presence of viral RNA, fresh tissues stored in diluent were homogenized for 1 min at 34 Hz with a TissueLyser II (Qiagen, Germantown, MD, USA), and 50 µL of homogenized material was used for viral RNA extraction utilizing the Mag-Bind DNA/RNA 96 kit (Omega Bio-tek, Inc., Norcross, GA, USA) and KingFisher Flex System with 96-deep well head (Thermo Fisher Scientific, Waltham, MA, USA). Extracted RNA was eluted in 50 µL of water.

After extraction, reverse transcription quantitative PCR (RT-qPCR) was conducted with 8 µL of RNA in a 20-µL reaction using the iTaq Universal One-Step RT-qPCR kit (Bio-Rad Laboratories, Hercules, CA, USA). Primers and probe targeting MADV nsP2, designed using IDT PrimerQuest, were: forward (5’-3’) – GGCTGAACAGGTGCTAGTTAT; reverse (5’-3’) – CTATTCCAATCCCGGACTTTCA; probe (5’-3’) – CGCGCCGGTAGGTACAAAGTAGAA (6-FAM/ZEN/3’ IB FQ). Amplicon size was 117 bp and final concentration of primers and probe were 400nM and 250nM, respectively. Reactions were run on a Bio-Rad CFX96 (Bio-Rad Laboratories, Hercules, CA, USA) under the following conditions: 50°C for 10 min, 95°C for 2 min; 40 cycles of 95°C for 15 sec and 60°C for 30 sec. Samples with cycle threshold (Ct) values ≤39 were considered positive. Using an in vitro-purified transcript, the limit of detection of the molecular assay was determined to be approximately 8 RNA copies per reaction.

Samples that were positive by RT-qPCR were subsequently tested by plaque assay to assess the presence and quantity of infectious virus. For the plaque assay, CCL-81 Vero cells were seeded in 6-well plates at 90-100% confluency. Sample dilutions were selected based on the RT-qPCR Ct value, with lower Ct values tested at higher dilutions. Aliquots (100 µL) of each sample dilution were inoculated onto the cell monolayers and allowed to adsorb for 1 h at 37°C in a 5% CO_2_ incubator with gentle rocking every 15 min. Following adsorption, 1 mL of 1% agarose overlay medium was added to each well. Plates were incubated for 48 h, after which a second agarose overlay containing 4% neutral red (1 mL/well) was added. Following an additional 24 h incubation at 37°C in the 5% CO_2_ incubator, plaques were counted and viral titers were calculated as plaque-forming units (PFU)/mL after accounting for the corresponding dilution factors.

### RNAemia

Because the volume of serum collected from mice prior to euthanasia was limited (5-15 µL), samples were brought to a final volume of 70 µL with DMEM supplemented with 5% FBS. Viral RNA was extracted from 50 µL of each diluted serum sample, and RNA extraction and RT-qPCR were performed as described for tissue samples.

### Histopathology

Tissue samples fixed in 10% neutral buffered formalin were routinely processed, paraffin embedded, sectioned, and stained with hematoxylin and eosin (H&E). For the brain, cross sections were trimmed at four representative anatomical levels following the protocol described by Rao et al. (26). Representative cervical, thoracic, and lumbar segments of the spinal cord were routinely decalcified prior to processing. Paraffin sections (4 µm) of collected tissues were examined blindly for microscopic lesions.

### Plaque reduction neutralization tests (PRNTs)

To determine anti-MADV neutralizing antibody titers, sera collected by cardiac puncture at the time of euthanasia were evaluated by PRNTs. CCL-81 Vero cells grown to 90-100% confluency in 6-well plates were used for all assays. Serum samples were heat-inactivated at 56°C for 30 min and mixed 1:1 with a 1:30,000 dilution of MADV-BR stock. The positive control consisted of a 1:1 mixture of 1x M199 medium (Thermo Fisher Scientific, Waltham, MA, USA) and the same 1:30,000 dilution of MADV-BR. The serum-virus mixtures were inoculated into the cell monolayers and allowed to adsorb for 1 h at 37°C in a 5% CO_2_ incubator. Following adsorption, 1 mL of 1% agarose overlay medium was added to each well. Plates were incubated for 48 h, after which a second agarose overlay containing 4% neutral red was added (1 mL/well). Following an additional 24 h incubation at 37°C in a 5% CO_2_ incubator, plaques were counted, and the percentage of plaque reduction was determined relative to the positive control. Serum samples were initially screened at a 1:10 dilution, and samples exhibiting ≥50% plaque reduction were subsequently tested at higher dilutions. Plaque reduction was calculated relative to the control (virus only). When plaque reduction crossed 50% within the dilution series, the PRNT_50_ titer was estimated by log10 interpolation between the two dilutions surrounding the 50% threshold. A PRNT_50_ titer of 10 was used as the cutoff for detectable neutralizing antibodies.

### Data analysis

For survival analysis, mice reaching study endpoints due to severe disease were considered nonsurvivors, whereas those euthanized for scheduled histologic analysis were censored at the time of euthanasia. Kaplan-Meier survival curves were compared among experimental groups using the log-rank test. Survival between females and males was subsequently compared within experimental groups that exhibited mortality using the same approach.

Body weight data recorded daily for 12 days after challenge were expressed as the percent change relative to day 0. Percent body weight change was analyzed using a linear mixed-effects model with experimental group, sex, dpi, and their interactions as fixed effects, with repeated measurements accounted for by mouse ID. Because sex and its interactions were not significant, the final model included only experimental group, dpi, and their interaction. When a significant group x dpi interaction was detected, pairwise comparisons among groups at each dpi were performed. Body weight at baseline was compared between females and males by *t*-test.

For visualization, the proportion of mice within each clinical severity category was calculated by experimental group at each dpi. Clinical scores among groups were compared using mixed-effects ordinal logistic regression with experimental group, dpi, and their interaction as fixed effects and mouse ID as a random intercept. When ordinal models failed to converge because of sparse data in one or more severity categories, clinical signs were analyzed as binary outcomes (present versus absent) using mixed-effects logistic regression. Posterior limb paralysis and convulsions were summarized descriptively because of their low frequency.

For tissue RT-qPCR data, only MADV-challenged groups (Groups 2 and 3) were included in the analysis. The proportion of RT-qPCR-positive samples was compared between groups separately for each tissue using Fisher’s exact test. Tissue samples positive for MADV RNA by RT-qPCR were further evaluated by plaque assay but because samples were not normalized by weight, plaque assay results were summarized descriptively only.

In the PRNTs, the proportion of mice with detectable MADV-neutralizing antibodies (PRNT_50_ ≥10) was compared among experimental groups using logistic regression adjusted for dpi, with Group 2 (Mock + MADV) as the reference group. For visualization purposes, PRNT_50_ titers were classified as negative (<10), low (10–40), intermediate (>40–<320), or high (≥320), and the number of mice within each category was plotted by experimental group and dpi.

Statistical analyses were performed using SAS 9.4 (SAS Institute Inc., Cary, NC, USA) through SAS Studio, and figures were generated using R version 4.6.1 (R Foundation for Statistical Computing, Vienna, Austria).

## RESULTS

### Prestige 3® vaccination protected mice from mortality following MADV challenge

Survival differed significantly among experimental groups (*p*=0.0075) (Fig. 2). All mice in Group 1 (Vaccine + Mock) and Group 3 (Vaccine + MADV) remained alive throughout the 30-day observation period. In Group 2 (Mock + MADV), 6 mice reached humane endpoints because of severe neurological disease between 6 and 15 dpi. Within Group 2, survival was significantly lower in females than in males (*p*=0.036) (Fig. 2).

**Fig. 2.**
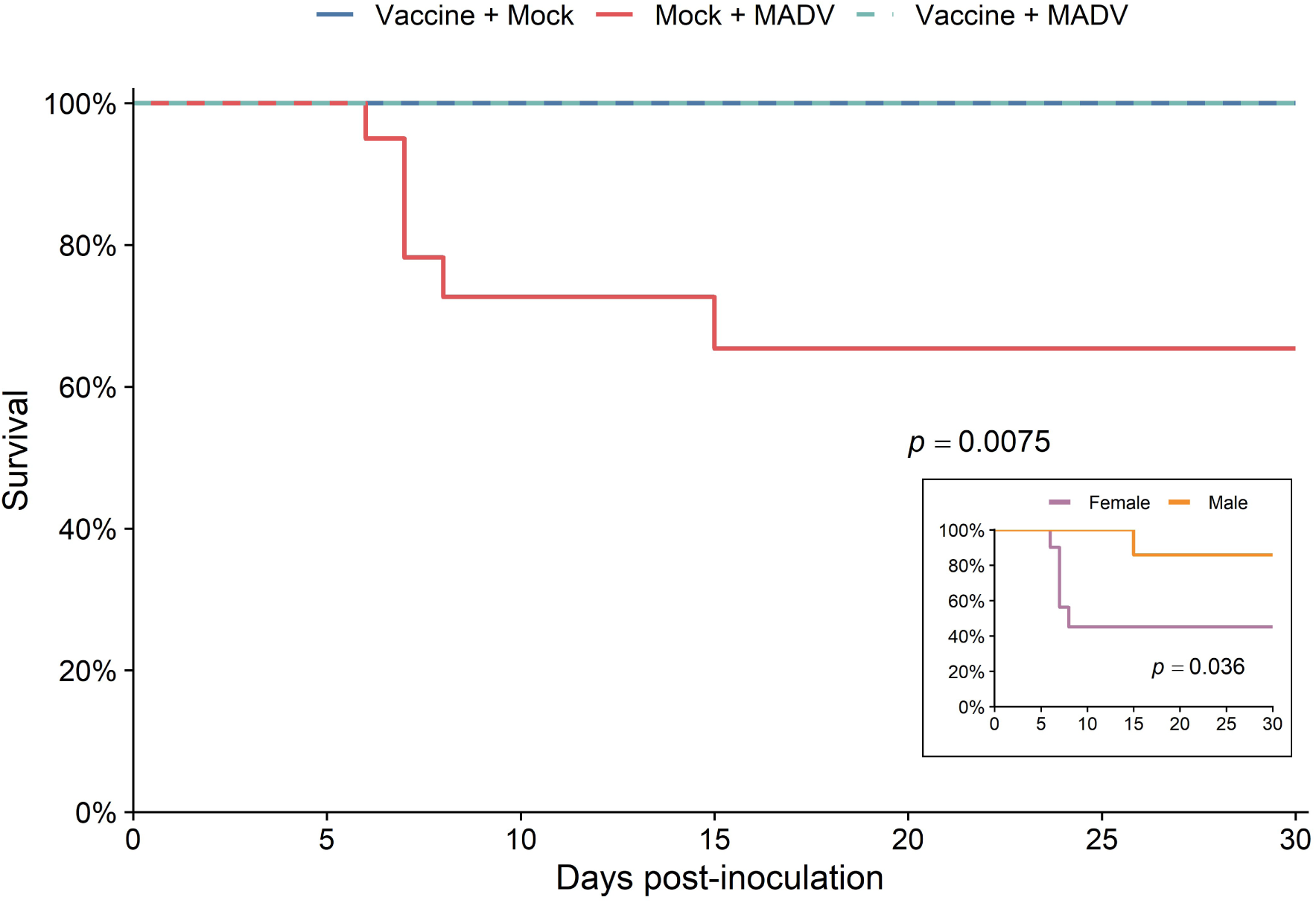
Survival of experimental mice. Mice were vaccinated with Prestige 3®, a trivalent vaccine containing inactivated North American eastern equine encephalitis virus (EEEV), and challenged with Madariaga virus (MADV), which belongs to the EEEV complex. Following challenge, survival was monitored daily for 30 days. Survival curves were generated using Kaplan-Meier and compared using the log-rank test. Experimental groups consisted of mice vaccinated with Prestige 3® and mock challenged (Group 1: Vaccine + Mock), mice mock vaccinated and challenged with MADV (Group 2: Mock + MADV), and mice vaccinated with Prestige 3® and challenged with MADV (Group 3: Vaccine + MADV). Survival differed significantly among groups (*p*=0.0075), with mortality occurring only in Group 2. Inset: Survival of female and male mice in Group 2, showing that females exhibited significantly higher mortality (*p*=0.036).

### Vaccination did not significantly prevent weight loss following MADV challenge

At baseline, males had significantly greater body weight than females (28.38 ± 1.67 g vs. 23.69 ± 1.28 g; *p*<0.0001). The pattern of body weight change over the 12-day period differed significantly among experimental groups (group x dpi interaction, *p*=0.0023) (Fig. 3). During the first 5 days after challenge, mice in Group 3 (Vaccine + MADV) lost more weight than mice in Groups 1 (Vaccine + Mock) (*p*<0.0001 per dpi) and 2 (Mock + MADV) (*p* ranging from <0.0001 to 0.0005 per dpi). From 6 dpi onward, both challenged groups (Groups 2 and 3) lost more weight than Group 1 (*p* ranging from 0.0053 to <0.0001 per dpi), whereas no significant differences were observed between Groups 2 and 3 during this period.

**Fig. 3.**
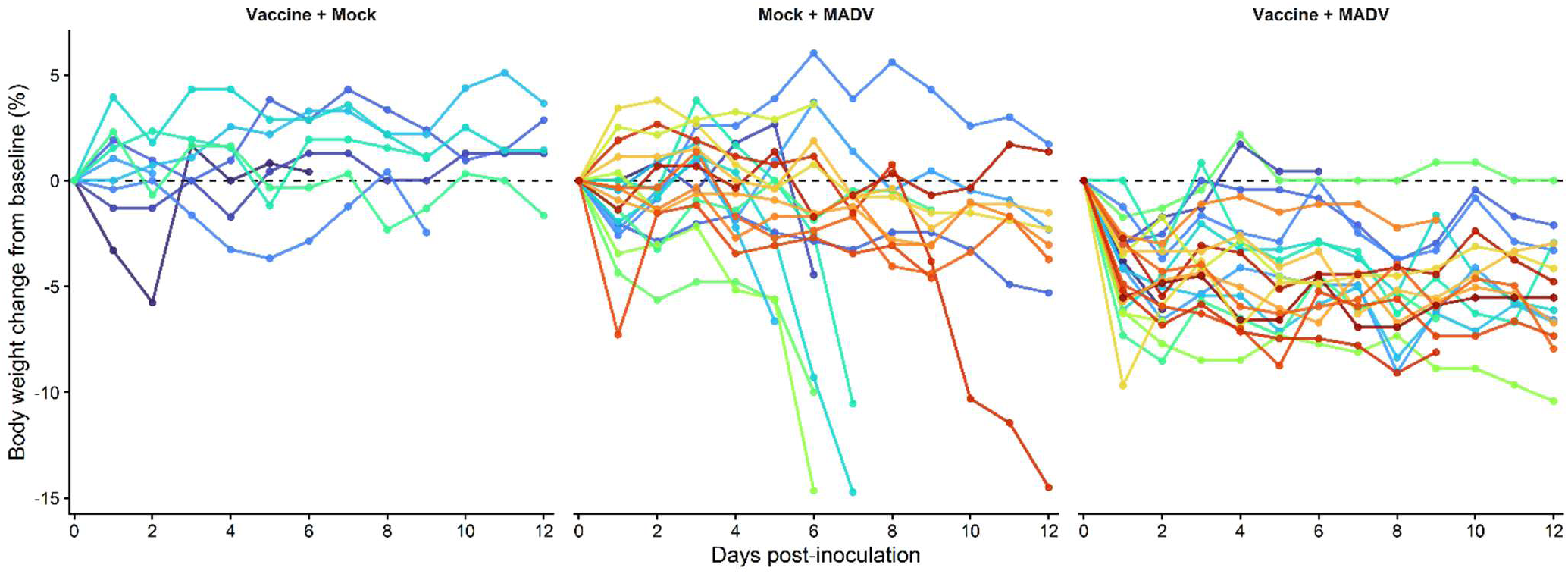
Change in body weight of experimental mice over time. Body weight was recorded daily in experimental mice over the 12-day clinical observation period. Groups consisted of mice vaccinated with Prestige 3® and mock challenged (Group1: Vaccine + Mock), mice mock vaccinated and challenged with Madariaga virus (MADV) (Group 2: Mock + MADV), and mice vaccinated with Prestige 3® and challenged with MADV (Group 3: Vaccine + MADV). During the first 5 days post-inoculation, Group 3 significantly lost more weight than Groups 1 and 2. From days 6 to 12, both challenged groups (Groups 2 and 3) lost more weight compared with Group 1.

### Vaccination reduced the frequency and severity of clinical signs following MADV challenge

The temporal distribution of clinical signs is shown in Fig. 4. Compared with mock-vaccinated mice challenged with MADV (Group 2), vaccinated mice challenged with MADV (Group 3) had significantly lower odds of developing orbital tightening, hunched back, and lethargy. The frequency of ruffled fur did not differ significantly between the challenged groups (Groups 2 and 3) (Table 1). Posterior limb paralysis and convulsions were observed in one mouse from Group 2. Conjunctivitis, which was not among the clinical signs monitored, was also observed in this mouse.

**Fig. 4.**
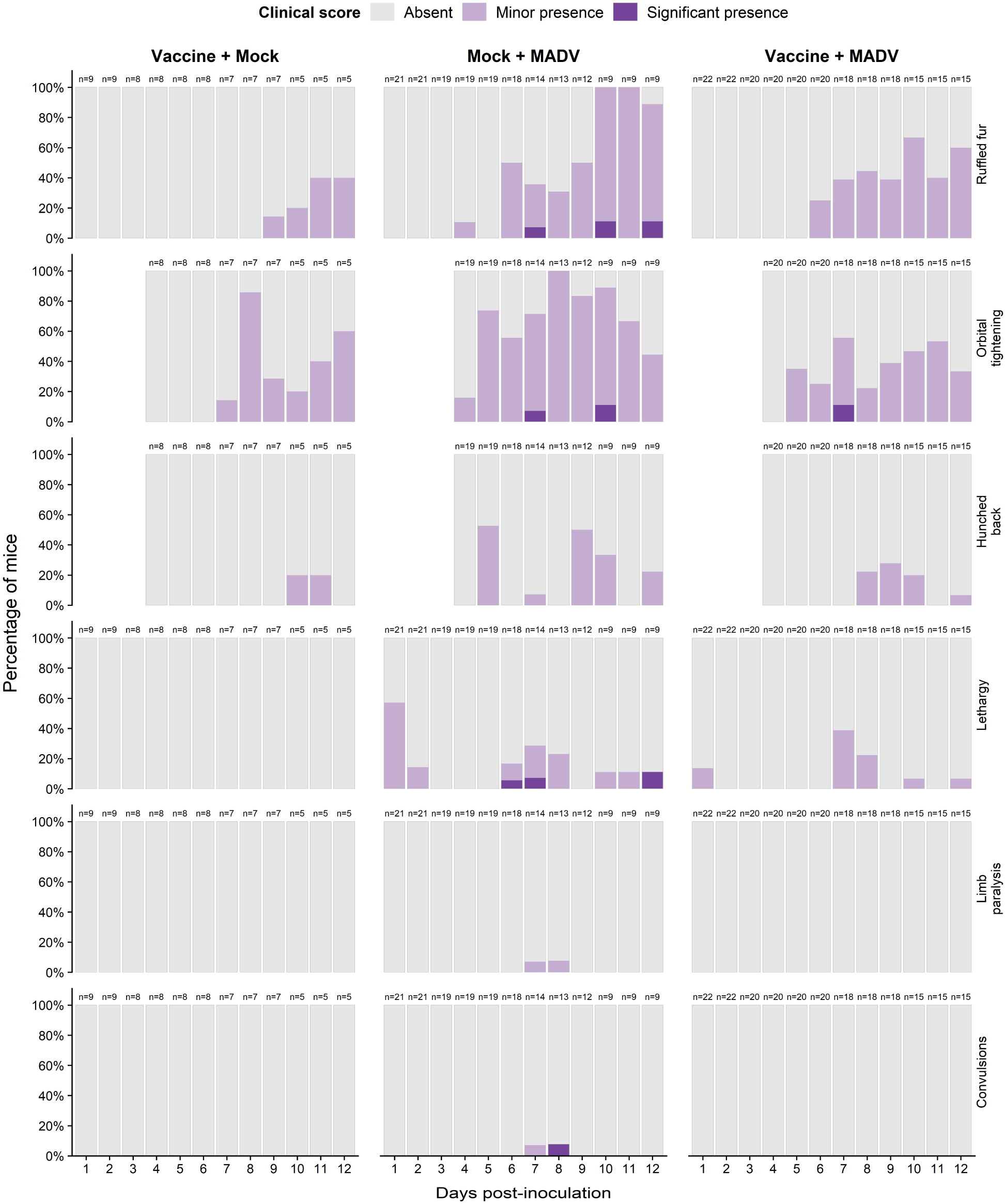
Clinical signs in experimental mice. Clinical signs (except body weight) were scored as absent, minor presence, or significant presence. The graphs show the percentage of mice exhibiting ruffled fur, orbital tightening, hunched back, lethargy, posterior limb paralysis, and convulsions at each day post-inoculation during the 12-day clinical observation period, stratified by clinical score. Experimental groups consisted of mice vaccinated with Prestige 3® and mock challenged (Group 1), mice mock vaccinated and challenged with Madariaga virus (MADV) (Group 2), and mice vaccinated with Prestige 3® and challenged with MADV (Group 3). Odds ratios derived from mixed-effects ordinal and logistic regression analyses are presented elsewhere. Overall, Groups 1 and 3 had significantly lower odds of exhibiting clinical signs and of developing more severe clinical signs than mock-vaccinated mice challenged with MADV (Group 2). Orbital tightening and hunched back were not recorded during the first three days post-inoculation.

**Table 1.** Comparison of clinical signs in experimental mice. Odds ratios (ORs), 95% confidence intervals (CIs), and *p*-values from mixed-effects models are shown.

| Clinical sign | Reference group | Comparison | OR (95% CI) | *p*-value |
| --- | --- | --- | --- | --- |
| Ruffled fur | 2 | Group 1 | 0.16 (0.06–0.46) | 0.0007 |
|  |  | Group 3 | 0.66 (0.37–1.20) | 0.176 |
| Orbital tightening | 2 | Group 1 | 0.26 (0.10–0.66) | 0.005 |
|  |  | Group 3 | 0.39 (0.20–0.76) | 0.006 |
| Hunched back | 2 | Group 1 | 0.15 (0.03–0.72) | 0.018 |
|  |  | Group 3 | 0.38 (0.17–0.86) | 0.021 |
| Lethargy | 2 | Group 3 | 0.33 (0.14–0.77) | 0.011 |

Clinical signs with adequate representation across all three severity categories were analyzed using cumulative logit mixed-effects models. Clinical signs with sparse observations in the highest severity category that resulted in non-convergence of the ordinal models were analyzed as binary outcomes (present versus absent) using mixed-effects logistic regression. Odds ratios are presented relative to Group 2 (Mock + MADV). Lethargy was compared between Groups 2 and 3 only, as no mice in Group 3 exhibited this clinical sign. Posterior limb paralysis and convulsions occurred in a single mouse and were not included in this analysis.

### Vaccinated mice had no detectable RNAemia but exhibited limited viral dissemination to tissues

No RNAemia was detected in vaccinated mice challenged with MADV (Group 3), whereas 15 of 22 mock-vaccinated mice challenged with MADV (Group 2) had at least one RT-qPCR-positive serum sample between 2 and 4 dpi (Fig. 5A). MADV RT-qPCR Ct values in positive serum samples ranged from 26.98 to 38.80.

**Fig. 5.**
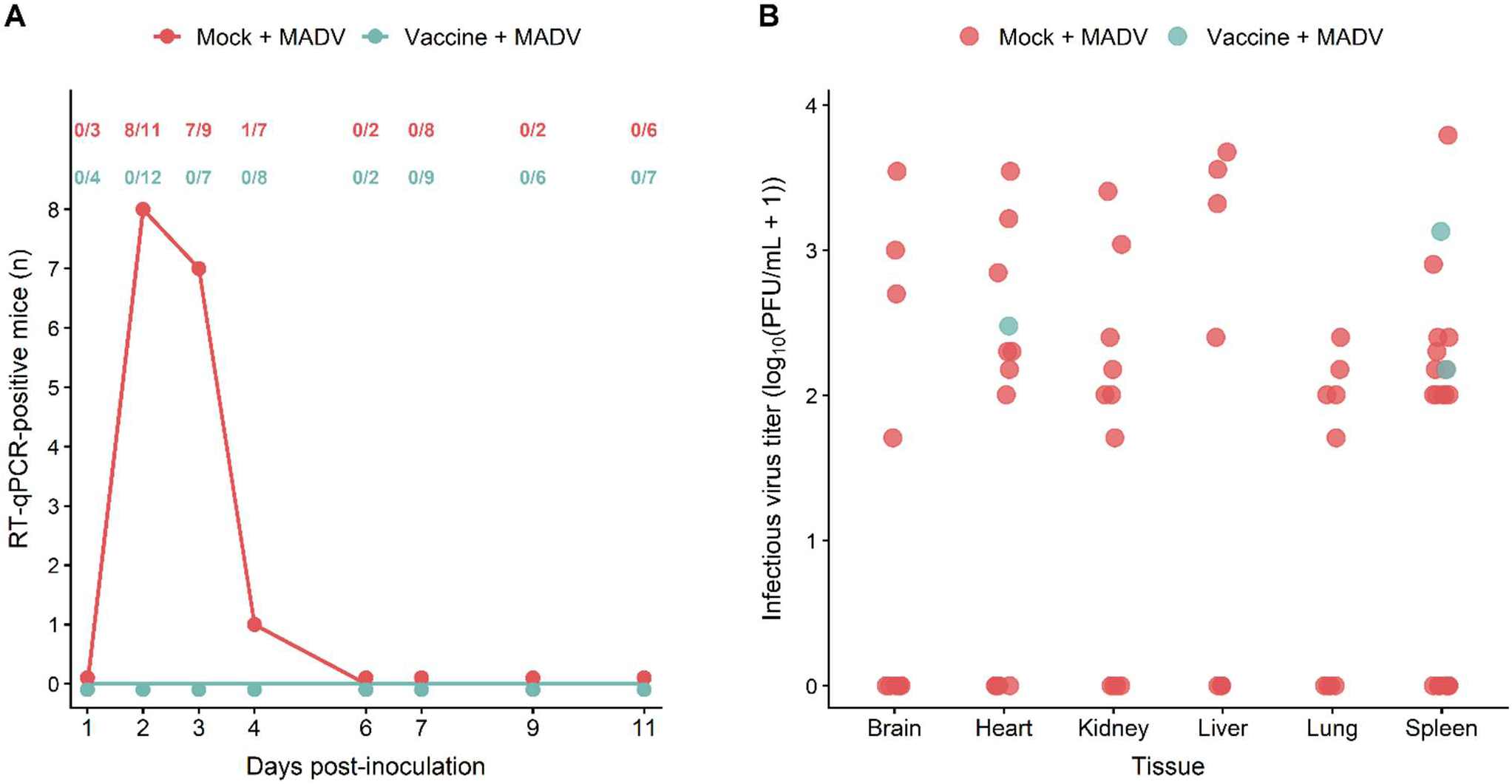
Serum RNAemia and infectious Madariaga virus (MADV) in tissues collected from experimental mice. (A) MADV RNA in serum from mock-vaccinated mice challenged with MADV (Group 2) and Prestige 3®-vaccinated mice challenged with MADV (Group 3) was assessed by reverse transcriptase quantitative PCR (RT-qPCR) during the 12-day clinical monitoring period. RNAemia was detected in Group 2 mice between 2 and 4 days post-inoculation but was not detected in Group 3 mice. (B) Tissue samples testing positive for MADV RNA by RT-qPCR were assayed by plaque assay. Infectious virus was recovered from multiple samples representing all tissue types in Group 2, whereas in Group 3 infectious virus was recovered from heart and spleen samples. Viral titers are presented as log_10_(PFU/mL + 1). RT-qPCR positivity at different dpi did not necessarily represent the same mice; 15 mice in Group 2 had at least one positive serum sample between 2 and 4 dpi.

Viral RNA was detected in all tissue types (brain, heart, kidney, liver, lung, and spleen) in Group 2 mice, whereas in Group 3 viral RNA was detected in heart and spleen samples. The frequency of MADV RNA detection was significantly lower in Group 3 compared with Group 2 for all tissue types (Table 2).

**Table 2.** Detection of Madariaga virus RNA in tissues of experimental mice by reverse transcriptase quantitative PCR (RE-qPCR).

| Tissue | Group 2<br>(Mock + MADV)<br>n/N<sup>1</sup> (%) | Group 3<br>(Vaccine + MADV)<br>n/N (%) | *p*-value<sup>2</sup> |
| --- | --- | --- | --- |
| Brain | 9/21 (42.9%) | 0/22 (0%) | 0.0005 |
| Heart | 11/21 (52.4%) | 1/22 (4.5%) | 0.0006 |
| Kidney | 9/21 (42.9%) | 0/22 (0%) | 0.0005 |
| Liver | 5/21 (23.8%) | 0/22 (0%) | 0.0211 |
| Lung | 8/21 (38.1%) | 0/22 (0%) | 0.0011 |
| Spleen | 20/21 (95.2%) | 0/22 (0%) | <0.0001 |

<sup>1</sup>Number of samples testing positive by RT-qPCR for Madariaga virus/total number of samples tested for that tissue. <sup>2</sup>*p* was calculated by Fisher's exact test.

Among RT-qPCR-positive samples, infectious virus was recovered from all tissue types in Group 2. Among plaque-positive samples, titers ranged from 5.0 × 10^1^ to 3.5 × 10^3^ PFU/mL in brain, 1.0 × 10^2^ to 3.5 × 10^3^ PFU/mL in heart, 5.0 × 10^1^ to 2.55 × 10^3^ PFU/mL in kidney, 2.5 × 10^2^ to 4.75 × 10^3^ PFU/mL in liver, 5.0 × 10^1^ to 2.5 × 10^2^ PFU/mL in lung, and 1.0 × 10^2^ to 6.2 × 10^3^ PFU/mL in spleen samples. In Group 3, infectious virus was recovered from heart (3.0 × 10^2^ PFU/mL) and spleen (1.5 × 10^2^ and 1.35 × 10^3^ PFU/mL) samples (Fig. 5B). Some brain samples yielding very low Ct values (10.38-13.82) by RT-qPCR did not yield plaques in the plaque assay. These samples were collected from mice that died of neurological disease or were euthanized for histopathological analysis and had CNS lesions observed on histopathology.

### Central nervous system (CNS) lesions were absent in vaccinated MADV-challenged mice

Histologic lesions in the brain and spinal cord were observed exclusively in Group 2 (Mock + MADV) mice. The lesions consisted of neutrophilic and lymphohistiocytic meningoencephalitis and poliomeningomyelitis with neuronal necrosis. Among the mice with CNS lesions were the nonsurviving mice (n = 6) and two mice euthanized at predetermined time points for histopathologic analysis, one at 2 dpi and one at 9 dpi. Representative images of CNS lesions in mice of group 2 are shown in Fig. 6.

**Fig 6.**
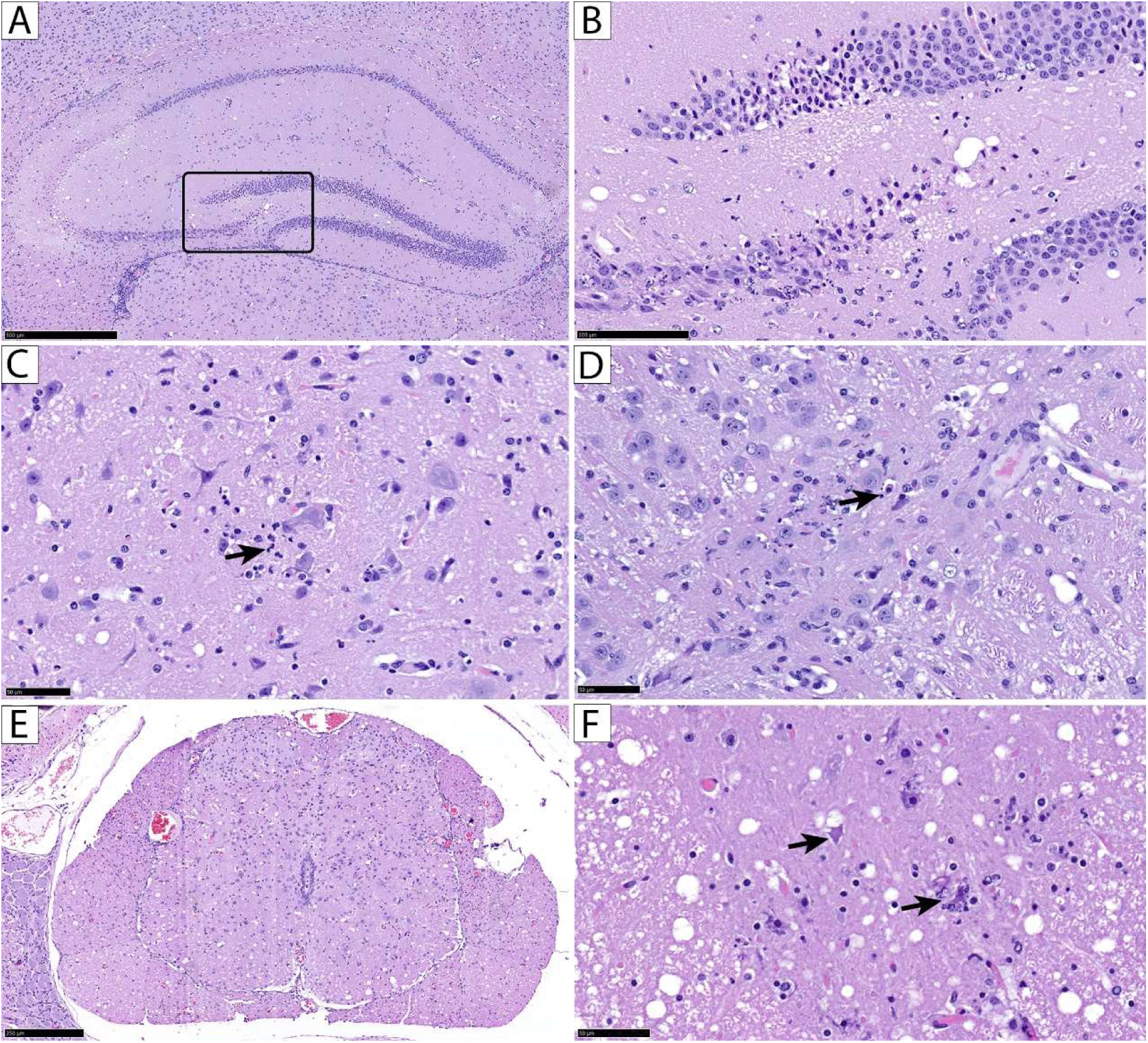
Histopathologic findings in the brain and spinal cord of Swiss mice infected with Madariaga virus, hematoxylin and eosin stain. (A) Hippocampus: locally extensive to multifocal neuronal necrosis of the pyramidal neuronal layer of CA1, CA2, CA3, and the granule cell layer of dentate gyrus, female mouse from Group 2. (B) Inset of A: Necrotic neurons are shrunken, triangular with hypereosinophilic cytoplasm, demonstrating acute neuronal necrosis. (C) Cerebrum: the neuropil shows infiltration of neutrophils (arrow) and glial cells, female mouse from Group 2. (D) Brainstem: multifocal neuronal necrosis (arrow) and gliosis, female mouse from Group 2. (E) Lumbar spinal cord: the grey matter of the spinal cord is hypercellular and infiltrated by inflammatory cells with mild lymphoplasmacytic meningitis, female mouse from Group 2. (F) Ventral horn of the thoracic spinal cord: Neuronal degeneration and necrosis, mouse female mouse from Group 2. Scale bars: (A), 500 µm; (B), 100 µm; (C), (D), and (F), 50 µm; and (E), 250 µm.

Neutrophilic and histiocytic infiltrates in the liver and kidneys were observed in mice from all experimental groups. Histiocytic capsulitis in the abdominal organs was observed only in vaccinated mice (Groups 1 and 3) and was associated with adjuvanct reaction. One mouse in Group 2 (Mock + MADV) and 2 mice in Group 3 (Vaccine + MADV) had neutrophilic vasculitis in the segmental artery of the paravertebral region. Focal myocarditis, focal alveolar hyperplasia in the lungs, hepatocellular microvesicular lipid accumulation, splenic lymphocytolysis, and interstitial nephritis with glomerulosclerosis and tubular atrophy were randomly and inconsistently observed. Detailed histologic findings for individual mice are summarized in Table S1.

### Protection in vaccinated mice occurred despite negligible MADV-neutralizing antibody titers

In Group 2, 16/18 (88.9%) mice had detectable MADV-neutralizing antibodies (PRNT_50_ ≥10), compared with 3/9 (33.3%) mice in Group 1 and 6/22 (27.3%) mice in Group 3. Because sera were collected at different dpi, differences among groups were evaluated using logistic regression adjusted for dpi. Experimental group was significantly associated with the presence of detectable neutralizing antibodies (p=0.0010), with lower odds in Group 3 (OR=0.007, p=0.0003) and Group 1 (OR=0.013, p=0.0030) compared with Group 2. When PRNT_50_ titers were categorized as negative (<10), low (10–40), intermediate (>40–<320), or high (≥320), all mice in Groups 1 and 3 had titers within the negative or low categories, including at later time points, whereas 15/16 Group 2 mice with detectable neutralizing antibodies had intermediate or high titers from 6–30 dpi (Fig. 7). Individual PRNT_50_ results are shown in Table S2.

**Fig. 7.**
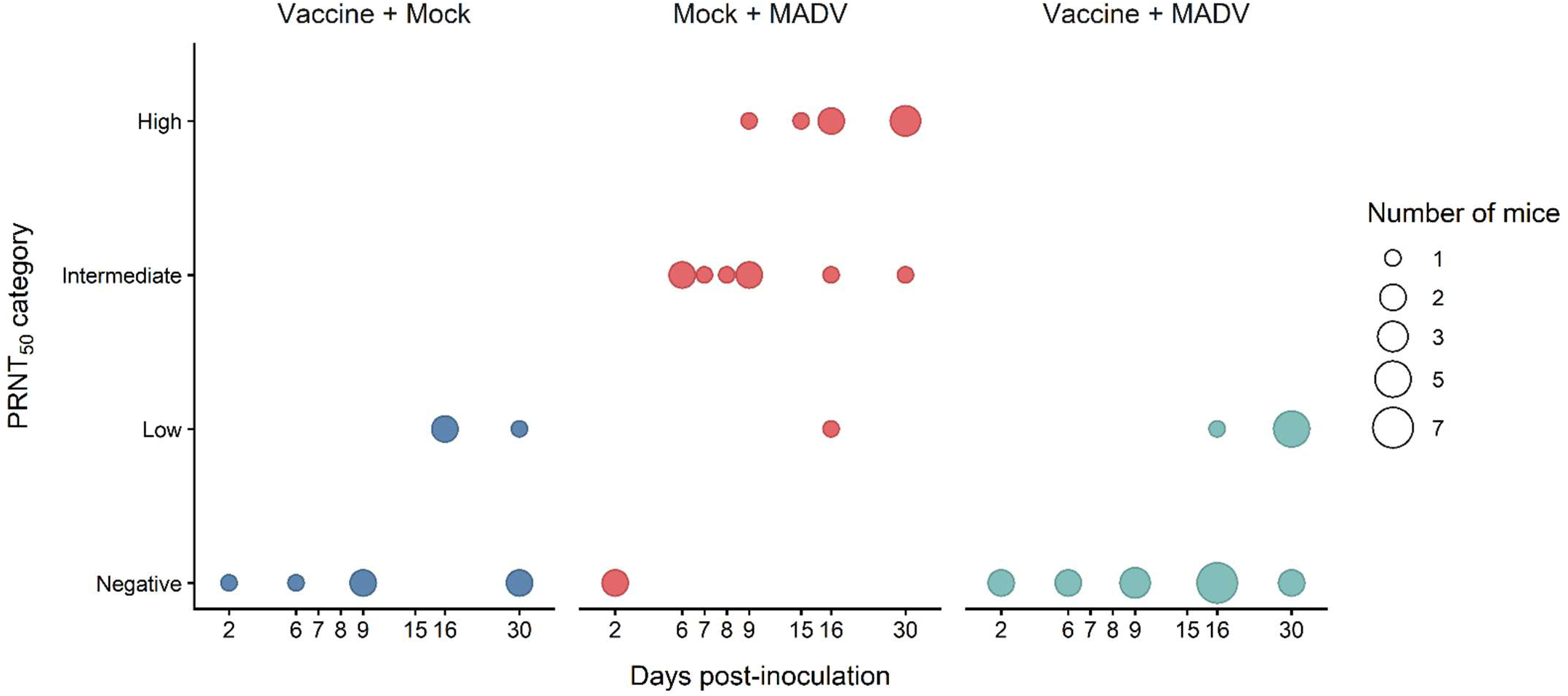
Neutralizing antibodies against Madariaga virus (MADV) in sera from experimental mice. Plaque reduction neutralization tests (PRNTs) were performed using terminal sera collected at necropsy (2-30 days post-inoculation). The reciprocal serum dilution resulting in a 50% reduction in plaques compared with virus-only controls (PRNT_50_) was calculated, and PRNT_50_ titers were categorized as negative (<10), low (10–40), intermediate (>40–<320), or high (≥320) for visualization purposes.

## DISCUSSION

Our findings in a murine model corroborate previous studies in humans and equids showing limited MADV-neutralizing antibody responses following NA EEEV vaccination (16, 20). However, whether these low titers translated into limited protection against MADV had not been evaluated. Our results show that vaccination with a trivalent inactivated NA EEEV vaccine can protect against severe disease and death following MADV infection despite negligible MADV-neutralizing antibody titers in mice. Vaccinated mice had PRNT_50_ titers <10 or in the low range (1:10–1:40) even at later time points (16–30 dpi), whereas nearly all unvaccinated mice sampled after 2 dpi developed intermediate (>1:40–<1:320) or high (≥1:320) titers. Still, none of the vaccinated, challenged mice died, whereas six unvaccinated mice reached study endpoints due to disease. Vaccination also reduced the frequency and severity of various clinical signs, except for ruffled fur, a nonspecific sign of distress.

Virological findings further supported the protective effect of vaccination. MADV RNA was not detected in the blood of vaccinated mice, whereas 18 of 22 unvaccinated mice had detectable RNAemia. Nevertheless, MADV RNA and infectious virus were detected in two spleen samples and one heart sample from vaccinated mice, indicating that vaccination did not completely prevent viral dissemination. Thus, vaccinated mice may have developed lower and/or shorter RNAemia that escaped detection because of the alternate-day blood collection schedule and/or assay sensitivity.

Histologic findings were also consistent with vaccine-mediated protection, as CNS lesions were observed only in unvaccinated, challenged mice. The CNS lesions consisted primarily of lymphohistiocytic and neutrophilic meningoencephalitis and poliomeningomyelitis with neuronal necrosis and neuronophagia. Similar CNS pathology has been reported in horses naturally infected with MADV in Brazil, including encephalitis or encephalomyelitis, neuronal necrosis and neuronophagia, meningitis, and spinal cord involvement (27–29). In the study by Silva et al., spinal cord lesions were generally mild, whereas those in our mice ranged from mild to marked. The CNS pathology observed in our mice also shares features with NA EEEV infection in humans (30) and experimental NA EEEV infection in C57BL/6 and CD-1 mice, in which neuronal tropism, neuronal necrosis, and neutrophilic encephalitis have been described (31, 32).

Histologic findings were also consistent with vaccine-mediated protection, as CNS lesions were observed only in unvaccinated, challenged mice. The CNS lesions consisted primarily of lymphohistiocytic and neutrophilic meningoencephalitis and poliomeningomyelitis with neuronal necrosis and neuronophagia. Similar CNS pathology has been reported in horses naturally infected with MADV in Brazil, including lymphoplasmacytic encephalomyelitis with neuronal death (29). The CNS pathology observed in our mice also shares features with NA EEEV infection in humans (30), horses (33) and experimental NA EEEV infection in C57BL/6 and CD-1 mice, in which neuronal tropism, neuronal necrosis, and neutrophilic encephalitis have been described (31, 32). Although vasculitis has been observed in human cases of NA EEEV (30), horses with MADV infection (29) as well as guinea pig and hamster models (31), this lesion was not a feature in NIH Swiss-infected mice.

An important finding of this study was that protection occurred despite limited MADV-neutralizing antibody responses. Although neutralizing antibodies are considered a major correlate of protection against viral infections and are widely used to evaluate vaccine-induced immunity, protection can also occur independently of neutralizing antibodies through T-cell responses and non-neutralizing antibody-mediated mechanisms (34–36). Much of the evidence for the protective role of non-neutralizing antibodies comes from studies using monoclonal antibodies targeting viral proteins. Protection can occur through several mechanisms, including Fc receptor-mediated mechanisms such as antibody-dependent cellular cytotoxicity (ADCC) and antibody-dependent cellular phagocytosis (ADCP), as well as complement-dependent cytotoxicity (CDC) (35, 36). A few studies have demonstrated the protective effect of non-neutralizing antibodies against alphavirus infections. In Earnest et al., non-neutralizing monoclonal antibodies against Mayaro virus protected mice from lethal infection through Fc-dependent mechanisms involving monocytes (35). An early study showed that non-neutralizing antibodies raised against Sindbis virus protected mice against homologous Sindbis virus challenge and also conferred protection against heterologous WEEV challenge (37).

The immune mechanisms responsible for protection in this study remain to be determined. An additional question is why vaccinated mice developed little to no MADV-neutralizing antibody response even after MADV challenge. One possibility is that pre-existing vaccine-induced immunity restricted viral replication, resulting in insufficient antigen exposure to induce a robust MADV-neutralizing antibody response. Alternatively, immunity against NA EEEV elicited by vaccination may have interfered with the development of a MADV-specific neutralizing antibody response following challenge (38).

A limitation of our work is the use of a trivalent vaccine (NA EEEV/WEEV/tetanus toxoid), as a commercial monovalent NA EEEV vaccine is not available, raising the possibility that immunity elicited by the WEEV component contributed to protection against MADV. Thus, although WEEV and NA EEEV have limited antigenic relatedness (39, 40), whether and to what extent the WEEV component contributed to the protection observed here cannot be determined from our data.

Our findings provide evidence of heterologous protection against MADV following vaccination with an inactivated equine NA EEEV-containing vaccine and support further investigation of the immune mechanisms responsible for this protection. The duration of both heterologous and homologous protection following vaccination is also an important question. In a previous study, horses vaccinated with an inactivated trivalent EEEV/WEEV/VEEV vaccine remained protected against NA EEEV challenge even when neutralizing antibodies were no longer detectable (40). Equine encephalitis virus vaccines are generally considered to provide relatively short-lived immunity, and annual booster vaccination is recommended (19). However, if waning immunity is inferred from declining neutralizing antibody titers, loss of detectable antibodies may not necessarily indicate loss of protection. Together, these observations raise questions about the extent to which neutralizing antibody titers alone reflect protective immunity following vaccination with inactivated equine encephalitis virus vaccines.

## DATA AVAILABILITY

Data supporting the findings of this study are provided in the supplementary material or are available from the corresponding author upon reasonable request.

## ETHICS APPROVAL

All procedures in this study were approved by the Texas A&M University Institutional Animal Care and Use Committee (protocol IACUC 2023-0269).

## ACKNOWLEDGMENTS

We thank the staff of the Global Health Research Complex (Biosafety Level 3 [BSL-3] facility) and the Biosafety Office at Texas A&M University (TAMU) for their assistance with optimization of BSL-3 protocols and execution of the experiments at the BSL-3 facility, and the staff at the TAMU Laboratory Animal Resources and Research facility for their assistance with animal care before challenge. We also thank Dr. Robert Rose and other personnel from the TAMU Comparative Medicine Program for assistance with mice protocols and consultation throughout the study. We are also grateful to Dr. Xiao Liang for assistance with the molecular assays.

This work was funded by TAMU AgriLife Research (FY24-25 Insect Vectored Diseases Seed grant to TM), and the United States Department of Agriculture National Institute of Food and Agriculture (Hatch # 6070 to TM). The funders had no role in study design, data collection and analysis, decision to publish, or preparation of the manuscript.

Contributions: Conceptualization: TM. Data Curation: SAC, TM, RRR, CS. Formal Analysis: SAC, TM, RRR, CS, WWT. Funding Acquisition: TM. Investigation: SAM, TM, RRR, CS, WWT. Methodology: TM, RRR, WT, GLH. Project Administration: TM. Resources: TM, RRR, GLH. Supervision: TM. Validation: SAC, TM, RRR, CS, WWT. Visualization, Writing – original draft: SAC, TM. Writing – review & editing: SAC, TM, RRR, GLH, WWT, CS.

